# Strong neutral sweeps occurring during a population contraction

**DOI:** 10.1101/2021.08.25.457712

**Authors:** Antoine Moinet, Stephan Peischl, Laurent Excoffier

**Affiliations:** Interfaculty Bioinformatics Unit, University of Bern, Baltzerstrasse 6, 3012 Bern, Switzerland; Swiss Institute of Bioinformatics, 1015 Lausanne, Switzerland; Institute of Ecology and Evolution, University of Bern, Baltzerstrasse 6, 3012 Bern, Switzerland

## Abstract

A strong reduction in diversity around a specific locus is often interpreted as a recent rapid fixation of a positively selected allele, a phenomenon called a selective sweep. Rapid fixation of neutral variants can however lead to similar reduction in local diversity, especially when the population experiences changes in population size, e.g., bottlenecks or range expansions. The fact that demographic processes can lead to signals of nucleotide diversity very similar to signals of selective sweeps is at the core of an ongoing discussion about the roles of demography and natural selection in shaping patterns of neutral variation. Here we quantitatively investigate the shape of such neutral valleys of diversity under a simple model of a single population size change, and we compare it to signals of a selective sweep. We analytically describe the expected shape of such “neutral sweeps” and show that selective sweep valleys of diversity are, for the same fixation time, wider than neutral valleys. On the other hand, it is always possible to parametrize our model to find a neutral valley that has the same width as a given selected valley. We apply our framework to the case of a putative selective sweep signal around the gene Quetzalcoatl in *D. melanogaster* and show that the valley of diversity in the vicinity of this gene is compatible with a short bottleneck scenario without selection. Our findings provide further insight in how simple demographic models can create valleys of genetic diversity that may falsely be attributed to positive selection.

## Introduction

Past demography and natural selection play a critical role in shaping extant genetic diversity. A central question in population genetics is to quantify their respective impact on observed genomic diversity. Because selection interferes with demographic estimates and vice versa, estimation of one of these two components is difficult without accounting for the other (Charlesworth *et al*. 1993, 1995; Kaiser and Charlesworth 2009; O’Fallon *et al*. 2010; Charlesworth 2013; Nicolaisen and Desai 2013; Johri *et al*. 2020, 2021b). Moreover, the relative importance of demography and selection as determinants of genome wide diversity is currently hotly debated, and may vary extensively among species (Corbett-Detig *et al*. 2015; Rousselle *et al*. 2018; Pouyet and Gilbert 2019; Galtier and Rousselle 2020). It has been shown that selection and demography can leave very similar footprints on the genetic diversity of a population (Andolfatto and Przeworski 2000; Teshima *et al*. 2006; Thornton and Jensen 2007; Johri *et al*. 2021a). Disentangling the effects of demography and selection is therefore crucial to avoid erroneous inference of evolutionary scenarios from genomic data (Jensen *et al*. 2005; Wares 2009; Mathew and Jensen 2015; Johri *et al*. 2020).

Hard selective sweeps lead to valleys of strongly reduced diversity around positively selected sites due to the hitchhiking of linked neutral loci (Maynard Smith and Haigh 1974), such observations of strong depletions of diversity in some genomic regions are often interpreted as due to past episode of positive selection, because the probability to observe a fast fixation of a neutral variant in a population of constant size is extremely low. However, during a range expansion for instance, some neutral or even mildly deleterious mutations can go quickly to fixation due to the low effective size of populations on the front of the range (Edmonds *et al*. 2004; Klopfstein *et al*. 2006; Hallatschek and Nelson 2008; Peischl *et al*. 2013), a phenomenon termed allele surfing (Klopfstein *et al*. 2006). Theoretical studies have shown that the average neutral diversity on the wave front decays exponentially as the range expands (Hallatschek and Nelson 2008), similarly to what happens when a population experiences a sudden decay of the population size, i.e. a population contraction, due to a drastic change in the environment for example. In both cases, a mutation appearing when the population size is shrinking might go quickly to fixation, inducing a strong decrease of diversity in the surrounding genomic region, whereas the average level of diversity might stay quite high depending on the strength and the duration of the contraction. As a result, the coalescent tree of alleles sampled in a population with strongly reduced effective population size will have short external branches, and long internal branches, depending on the parameters of the model (Excoffier *et al*. 2009). The site frequency spectrum associated to such a tree resembles a neutral SFS, but with a lack of rare alleles and an excess of high frequency sites, i.e. it becomes “flatter” (Sousa *et al*. 2014; Peischl and Excoffier 2015). The footprint left by the rapid fixation of a neutral allele on the surrounding genomic diversity, might thus be like that of a positively selected allele sweeping through a constant size population.

The expected shape of nucleotide diversity in genomic regions surrounding a site undergoing a rapid neutral fixation has been investigated analytically and numerically. Tajima (1990) studied the reduction of diversity during a neutral fixation at a given recombination distance from the fixing site. His results rely on rigorous mathematical arguments based on diffusion theory, but no closed form solution is provided for the shape of a neutral sweep. Johri *et al*. (2021a) described the valley of diversity occurring around a neutral fixation using an approach introduced for selective sweeps, assuming that the evolution of the allele frequency is that of a selected allele except in the initial stochastic phase. Here, we extend this work by inferring the dynamics of fixation of neutral alleles after a population contraction and we examine their effects on neighboring regions of the genome. We provide an analytical result for the expected coalescence time as a function of the recombination distance from the locus undergoing a fast fixation. Importantly, our results apply regardless of the process driving the allele going to fixation (neutrality, positive selection, background selection), as it only relies on the typical trajectory of an allele going to fixation in a given time, even though this trajectory differs depending on the underlying driver of this fixation (i.e., neutrality or selection). We compare our results against simulations and find that they hold for a wide range of realistic parameter combinations. We compare our results about the signature of neutral sweeps to patterns expected under selective sweeps and discuss potential differences between the signatures that could potentially allow us to discriminate between neutral and selective processes for a given demographic scenario. Finally, we investigate the similarity between the genomic signature of an allele going to fixation either selectively or neutrally and observe that a selective sweep signal can in principle be replicated in a neutral model with an appropriate choice of demographic parameters. To illustrate this point, we examine a classical example of a selective sweep found in the genome of *D. melanogaster* around the Qtzl gene (Rogers *et al*. 2010). We conclude that strong diversity depletions in the genome of a population, often attributed to the effect of positive selection, can be obtained with demographic effects only, and we call for caution when trying to detect signals of adaptation from genomic data, adding support to previous studies reaching similar conclusions (Thornton and Jensen 2007; Crisci *et al*. 2013; Jensen *et al*. 2019).

## Model

We model here the effect of an instantaneous population contraction on genomic diversity. Throughout the whole manuscript, time is measured backwards. We assume that *t*_c_ generations before the present, the population size instantaneously dropped from *N_0_* diploid individuals to *N_c_* individuals with *N_c_* < *N_0_*. We assume a standard coalescent model (Kingman 1982a; b) with discrete non-overlapping generations, random mating, monoecious individuals, and no selection. Two haplotypes sampled in the current population at time *t* = 0 have, as we go backwards in time, a constant probability (2*N_c_*)^-1^ of coalescing at each generation, for the first *t*_c_ generations, and then this probability switches to (2*N*_0_)^-1^ as we enter the ancestral uncontracted population. We can approximate the distribution of coalescence time *T* of these two haplotypes as a piecewise exponential distribution (see Appendix) with expected value:

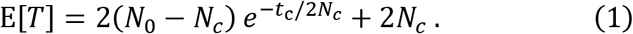

We see that the expected coalescence time decreases exponentially with the age of the contraction *t*_c_ and that it approaches 2*N_c_* for a very old contraction. Coalescence times cannot be measured directly from empirical data, but they are closely related to nucleotide diversity *π*. Under the infinitely many sites model, the number of nucleotide differences between two homologous DNA segments is proportional to their coalescence time *T* as *π* = 2*μT*, where *μ* is the total mutation rate for the whole segment. Multiplying eq. (1) by 2*μ* shows that an instantaneous population contraction leads to an exponential decrease of the expected nucleotide diversity along the genome with the age of the contraction *t*_c_. However, it does not inform us on the distribution of nucleotide diversity *π* along the genome, or on spatially correlated patterns of diversity such as local depletion or excess of diversity relative to the expectation.

Fig. 1 shows the evolution of the distribution of *π* as a function of the time *t*_c_ elapsed since the contraction. For *t*_c_ = 0, there is no contraction, and the population size remains constant and equal to *N*_0_. In this case we see (fig. 1a, 1b, *t_c_* = 0) that the distribution of π is symmetric and centered at E[π] = 4*N*_0_*μ*. For an older contraction, we see that the distribution is not only shifted to lower values of diversity as expected from eq. (1), but that it also becomes strongly peaked around *π* = 4*N*_c_*μ*. This bimodality of the distribution can be understood intuitively in the following way. There are two possible types of coalescent trees for haplotypes sampled after the population contraction (note that the tree depends on the locus considered because of recombination). Indeed, the most recent common ancestor (MRCA) of the sample lived either before the contraction (*T*_MRCA_ > *t_c_*), or after the contraction (*T*_MRCA_ < *t_c_*). In the former case, the tree at this locus has long inner branches and short outer branches, whereas in the latter case, the tree is essentially a (short) neutral tree corresponding to a population of constant size *N*_c_ (Excoffier *et al*. 2009). Both types of trees occur at different loci and correspond to the two observed modes in the distribution of the nucleotide diversity along the chromosome. The precise shape of the distribution of nucleotide diversity across sites depends on the relative frequency of both types of trees, which itself depends on the age of the contraction *t_c_*. For a sample of size two, the probability that the MRCA lived after the contraction, that is, *T*_MRCA_ < *t_c_* is 1 – *e*^−*t_c_*/2*N_c_*^. For a larger sample of haplotypes, there is no closed form solution for this probability, but the trees rooted after the contraction are rare for *t_c_* ≪ 2*N*_c_ and very frequent when *t_c_* ≫ 2*N*_c_ (Tavaré 1984). Therefore, the evolution of the distribution of *π* for increasing contraction age *t_c_* appears to be a transition from a unimodal distribution centered at 4*N*_0_*μ* to another unimodal distribution centered at 4*N*_c_*μ*, with both modes coexisting for intermediate ages (fig. 1). This bimodality has been pointed out previously in the context of population bottlenecks (Austerlitz *et al*. 1997); however, those studies mainly focused on long duration bottlenecks (the effect of a contraction or a bottleneck on nucleotide diversity is the same provided that the bottleneck is not yet finished, or that it finished very recently so that the effect of population recovery is negligible). In the present work, we investigate the effect of short contractions on the genetic diversity and make the claim that this short contraction regime is of particular interest as it can lead, such as in fig. 1c, to genomic signatures similar to those generated by positive selection acting on a few sites in an otherwise neutral genome. More specifically, we want to quantitatively describe the reduction of diversity along the genome that is observed around a locus with a small *T*_MRCA_ (such as in fig. 1c in the regions around 10-11 and 19-20 Mb), where we observe a valley or trough of diversity. Akin to what is done for selective sweeps, we consider the (neutral) fast fixation of an allele and analyze the impact of hitchhiking on the genetic diversity of neighboring loci, and we refer to this process as a neutral sweep.

**Figure 1.**
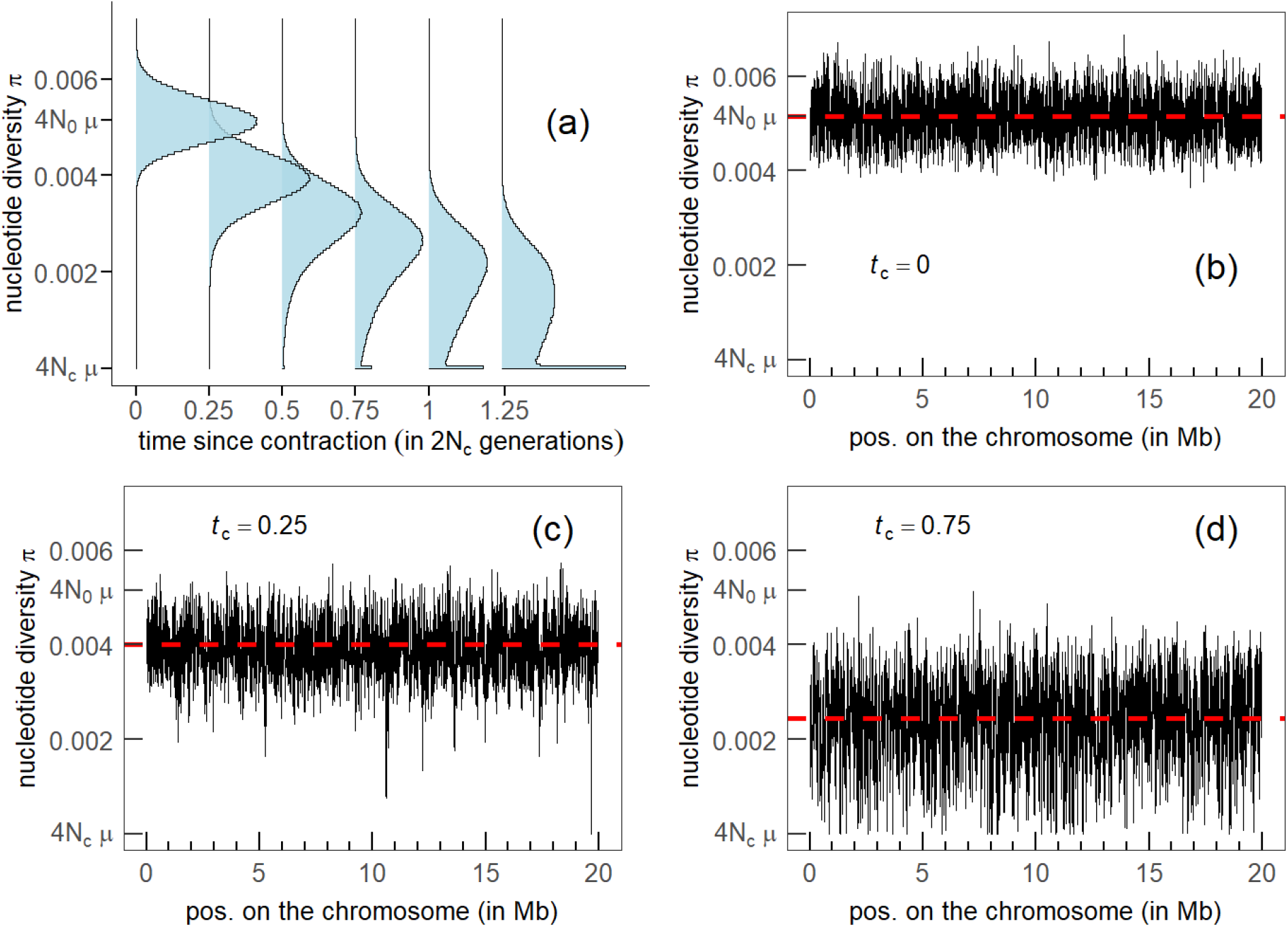
*Nucleotide diversity of a population experiencing a contraction, as a function of the time tc elapsed since the contraction, measured in units of 2Nc. (a) distribution of nucleotide diversity as a function of time, nucleotide diversity along the chromosome at t_c_ = 0 (panel b), at t_c_ = 0.25 (panel c) and at t_c_ = 0.75 (panel d). Population size before contraction N_0_ = 2.37×10^6^ and after contraction N_c_ = 4,400. Mutation rate μ = 5.42×10^-10^ per site per generation. Recombination rate r = 3.5×10^-8^ per site per generation. Chromosome size L = 20 Mb. Window size 10 Kb sliding at 1 Kb intervals. Sample size: 30 haplotypes. These parameters are taken from* Rogers *et al*. (2010). *Simulations were performed with fastsimcoal2* (Excofffier *et al*. 2021).

To investigate neutral sweeps in our model, we consider the following scenario: *t*m generations ago a mutation occurred at a single site on the chromosome, which we call the focal site. We further assume that this mutation has just fixed in the population, i.e., that it was segregating at a frequency strictly lower than one in the last generation (at *t* = 1), and has now (at *t* = 0) a frequency equal to one. We assume that the population contraction occurred *tc* generations ago, with *t_c_* ≥ *t*_m_. As the mutant enters the population as a single allelic copy at the focal locus, defined as a non-recombining region surrounding the focal site, this copy is a common ancestor for all the copies (2*N*_c_) present at fixation. However, it is not necessarily the most recent common ancestor. Fig.2 shows a sketch of our model to help visualize how recombination can maintain diversity at linked loci around a locus where a new mutation quickly fixed in the population.

**Figure 2.**
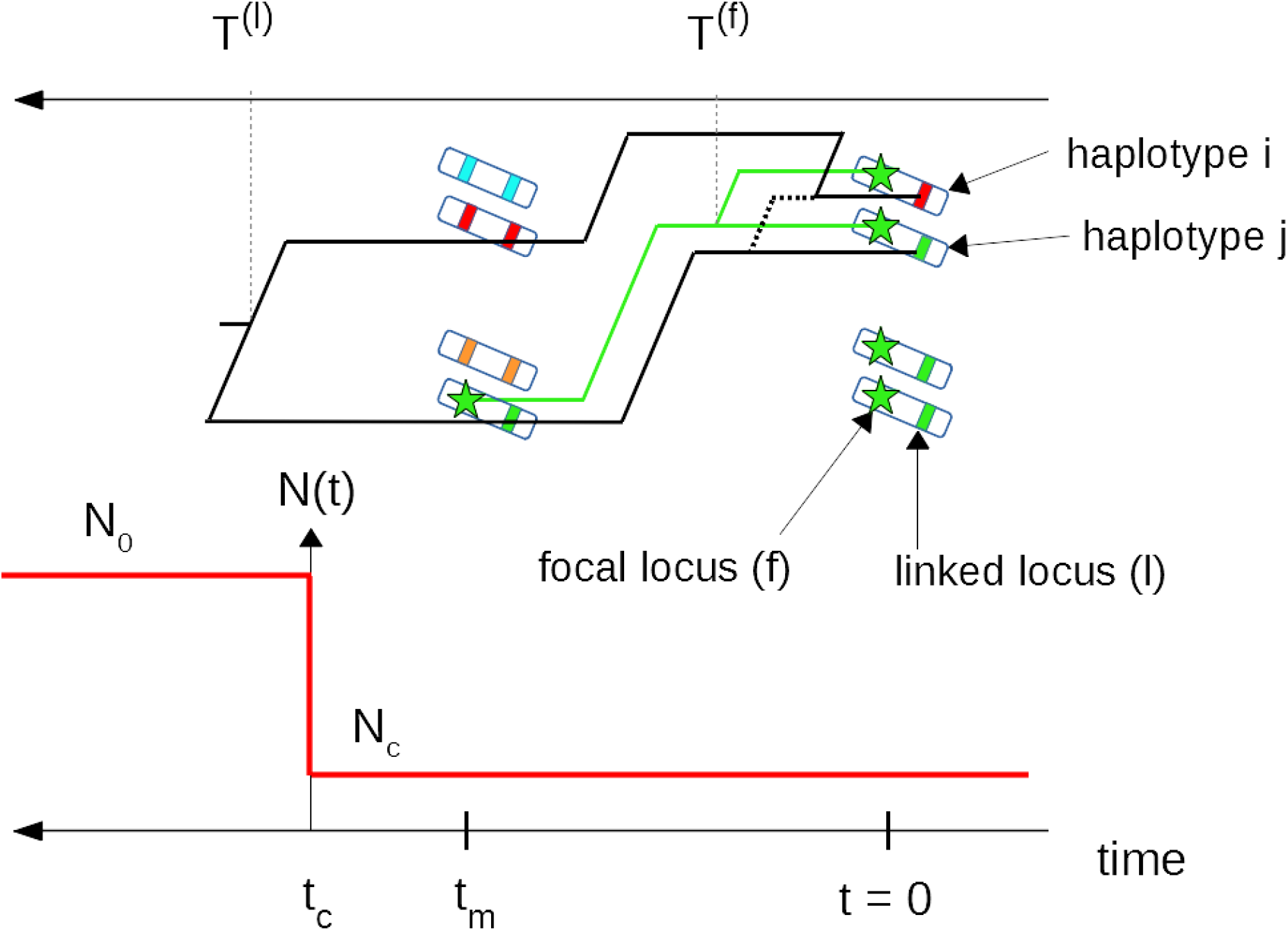
Instantaneous population contraction with a subsequent neutral fixation. A mutant (green star) appeared tm generations ago and has just fixed neutrally in a diploid population that experienced a contraction tc generations ago. We represent the population as a set of 2N_c_ two-locus haplotypes that are painted so that the gene copies present at t = 0 can be traced back to t = t_m_. Due to recombination, haplotype i carries a red gene copy at the linked locus at t = 0. Correspondingly, the coalescence time T^(l)^ of the haplotypes i and j at the linked locus (black tree) is larger than t_m_. On the other hand, the coalescence time T^(f)^ at the focal locus (green tree) is smaller than tm because at this locus all gene copies descend from the same haplotype (due to the fixation of the focal mutation).

## Results

### Average coalescence time at a linked locus

We can calculate the expected coalescence time *T*^(l)^ of two randomly sampled haplotypes at a linked locus as a function of the recombination rate *r* from the focal locus. The idea is to consider two haplotypes with a given coalescence time *T*^(f)^ at the focal locus, and then follow the genealogy of the gene copies carried by these two haplotypes at the linked locus backward in time, while considering possible recombination events. The expected coalescent time at the linked locus is then

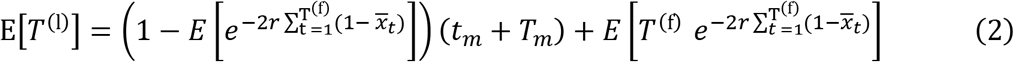

where 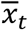 is the average frequency of the mutant (derived) allele at the focal locus at time *t* counting backward from present. A detailed derivation of this equation is given in Appendix A4. The first term of the right-hand side of eq. (2) corresponds to cases where lineages escape the neutral sweep due to recombination, and still have not coalesced after *t*_m_ generations. In this case we need to wait on average *T_m_* = 2(*N*_0_ – *N_c_*) *e*^−(*t_c_* – *t_m_*)/2*N_c_*^ + 2*N_c_* extra generations before the lineages coalesce, due to the contraction that happened *t*_c_ - *t*_m_ generations before the focal mutation. The second term of the right-hand side of eq. (2) corresponds to cases where the lineages cannot escape the sweep and are forced to coalesce at a time *T*^(l)^ ≤ *t*_m_.

### Distribution of coalescence times at the focal locus

To evaluate eq. (2), we need to determine the probability distribution of the pairwise coalescence times *T*^(f)^ at the focal locus, as well as the expected frequency trajectory of the derived allele. Even though this allele fixes neutrally in a population of constant size (the contraction occurs prior to the mutation), the distribution of coalescent times at the focal locus *T*^(f)^ departs from the usual exponential distribution for a neutral coalescent process because the allele fixes in exactly *t*_m_ generations, and hence the coalescence time for a randomly chosen pair of haplotypes is at most *t*_m_. Slatkin (1996) investigated the coalescent process within a “mutant allelic class” that originated from a single mutation at a given time in the past. He derived exact analytical results for the average pairwise coalescence time, but the coalescence distribution itself can only be expressed with multidimensional integrals and obtaining a closed form expression does not appear feasible. We therefore use a different approach: given a particular fixation trajectory of the mutant allele, i.e. given the number of mutant copies *N*_μ_ at each generation between *t* = 0 and *t* = *t*_m_, we can express the coalescence time-distribution within the mutant allelic class, using the result of a coalescent in a population with a time-dependent (but deterministic) size *N*_μ_(*t*) (Griffiths and Tavaré 1994). Averaging over all possible trajectories of the mutation, we obtain:

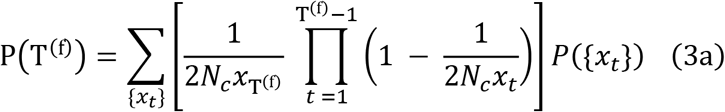

where *x_t_* = *N*_μ_(*t*)/(2*N_c_*) is the frequency of the mutant *t* generations from fixation, and *P*({*x_t_*}) is the probability of a given trajectory. *P*({*x_t_*}) can be evaluated (see Appendix A2) and the sum in eq. (3a) can in principle be computed numerically; however, the number of trajectories to consider is prohibitive. As a first approximation, we can replace *x_t_* by its expectation 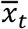, i.e., we neglect the fluctuations of the trajectory around the mean to obtain

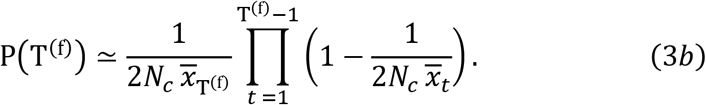

The last step is to determine the average trajectory of an allele fixing in exactly *t*_m_ generations. Zhao *et al*. (2013) as well as Maruyama and Kimura (Maruyama and Kimura 1975) have investigated the characteristic trajectory of an allele fixing in a given time but they do not provide a closed form solution. Here, we use a different approach (also based on diffusion theory to obtain an approximation for the average trajectory of an allele fixing in exactly *t*_m_ generations, starting from a frequency *p*_0_. As detailed in the Appendix A2, we obtain

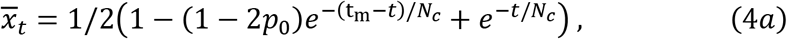

which is valid for *t*_m_ ≫ 2*N_C_*. For very fast fixations, i.e., when *t*_m_ ≪ 2*N_c_*, the frequency of the allele increases approximately linearly as

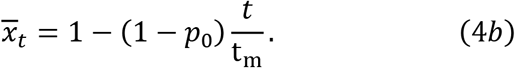

We remind the reader that *t* is counted backwards from fixation. Fig. 3 compares equations (4a) and (4b) to trajectories obtained from simulations of a Wright-Fisher diploid population. We find good agreement between the simulations and the analytical results. Importantly, the typical neutral trajectory for large values of the fixation time has an “inverse-sigmoid shape” (fig. 3c), contrary to the typical sigmoid trajectory of a positively selected allele going to fixation in a constant size population (see fig. 5a). This neutral trajectory occurs because, conditional on non-loss, neutral alleles need to quickly escape loss at the beginning and remain at intermediate frequencies to stay away from both fixation and loss until they eventually fix in the population at *t* = 0 (i.e. in exactly *t*_m_ generations). Fig. 3d–3f also shows the coalescence time distribution for several values of the fixation time *t*_m_. The comparison of the distribution of pairwise coalescence time with numerical simulations of a Wright-Fisher model shows that our approximation eq. (3b) is quite accurate but overestimates the probability of coalescence for large coalescence times when *t*_m_ is small (fig. 3d). Notably, coalescence (simulated or theoretical) is more probable at large times (i.e. when the mutant appeared) for short fixation times (fig. 3d), whereas it is more probable at small times (i.e. close to fixation) for large fixation times (fig. 3e). The coalescence rate within the mutant allelic class is given by the inverse of the number of mutant copies and is for all values of the fixation time slightly more than 1/2*N*_c_ at the first generation. However, when the fixation time is short (fig. 3d), there is a fast increase of the coalescence rate backwards in time, and many lineages are forced to coalesce at *t* = *t*_m_. When the fixation time is large (fig. 3f), the coalescence rate also increases backwards in time, but the increase is much slower. In that case, most coalescence events happen in much less than *t*_m_ generations, so that the early increase in frequency of the mutant has almost no influence on the coalescence distribution.

**Figure 3.**
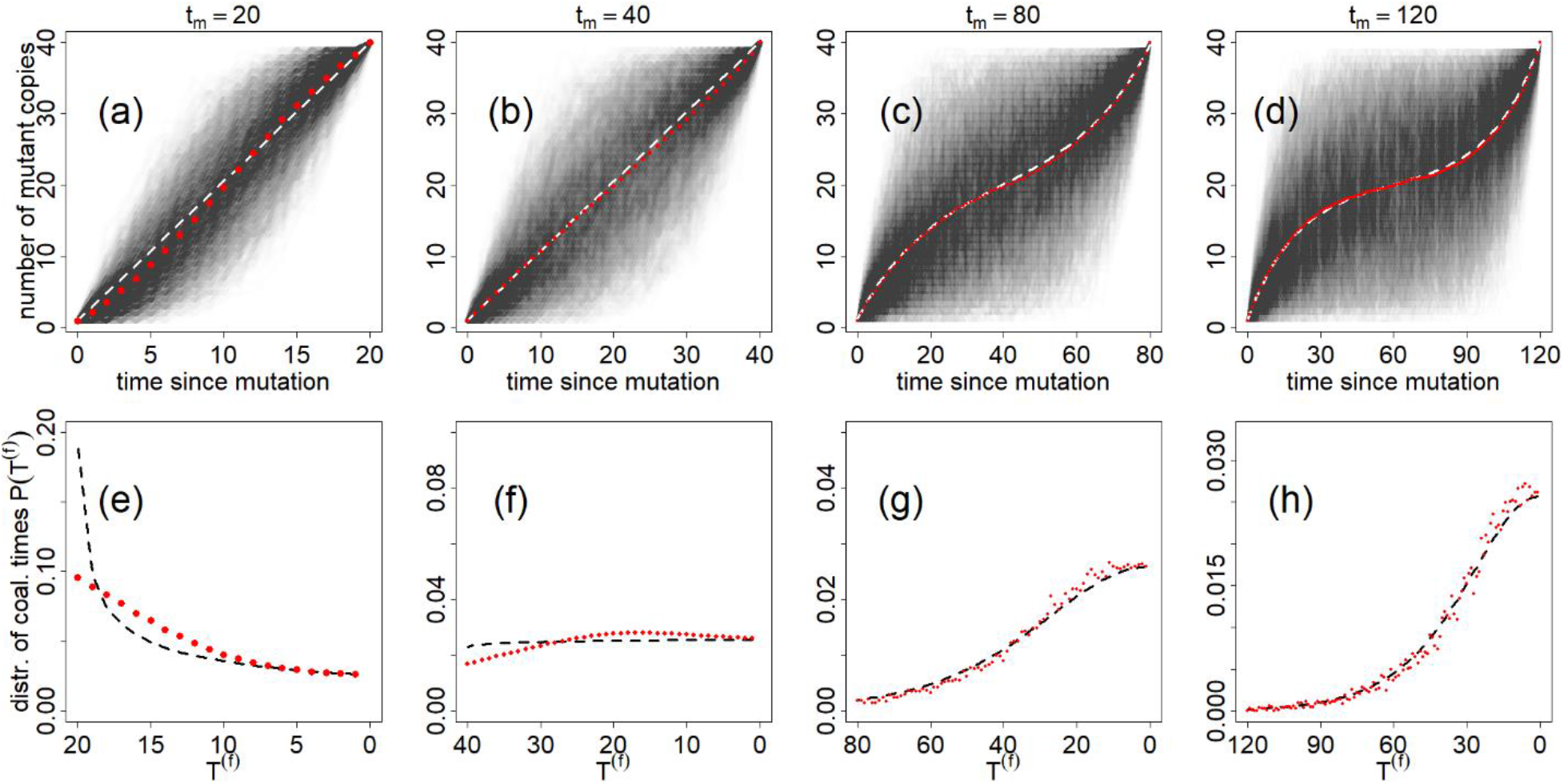
Average frequency (a-d) and coalescence time distribution (e-h) of an allele fixing in a diploid population of constant size N_c_ = 20 in exactly t_m_ generations, starting as a single copy (i.e. p_0_ = (2N_c_)^-1^). The red dots are the results of Wright-Fisher simulations, and the black and white dashed lines are calculated with eqs. (4b) (first and second columns) (4a) (third and fourth columns) and (3b). In panes (a-d) we show the variability of the fixation process by overlapping 1780 fixing trajectories. The (numerically estimated) probability, for a mutant that appears at the onset of the contraction, to fix in less than t_m_ generations is 0.006, 0.16, 0.64 and 0.86 for t_m_ = 20, 40, 80 and 120 respectively (for this particular value of Nc).

### Effect of a neutral sweep on linked diversity

Combining equations (3b), (4a) with eq. (2) allows us to get an approximation for the average coalescence time at linked loci. Since the derivation of eq. (2) assumes that there is at most one recombination event in the genealogy of a randomly chosen pair of gene copies, we expect it to be only accurate for small values of the recombination rate *r*. For large values of *r* we use a heuristic approach combining the result of eq. (2), which is accurate for small *r*, and the expected diversity at unlinked loci, which is equal to *T*_0_ = 2(*N*_0_ – *N_c_*) *e*^-*t_c_*/2*N_c_*^ + 2*N_c_* as stated in eq. (1). We fit the trough of diversity with an exponential function of the form:

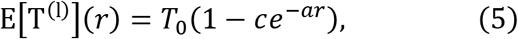

where the coefficients *c* = 1 – *E*[T^(f)^]/*T*_0_ and 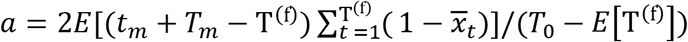 are obtained by imposing that eqs. (2) and (5) coincide for small values of *r* (using a linear expansion in *r*). On fig. 4 we compare the result of eq. (5) to Wright-Fisher simulations with two recombining loci. We see in fig. 4a that the exponential function fits the data accurately at large values of the recombination distance, but that the fit is biased for intermediate values of *r*. In fig. 4b we see that the approximation is very good for low values of the recombination distance, although there still is a slight bias. This discrepancy at small *r* can be corrected (solid lines in fig. 4) if we use numerical estimations of 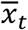 and P(T^(f)^), instead of eqs. (4) and (3b), to evaluate eq. (5).

**Figure 4.**
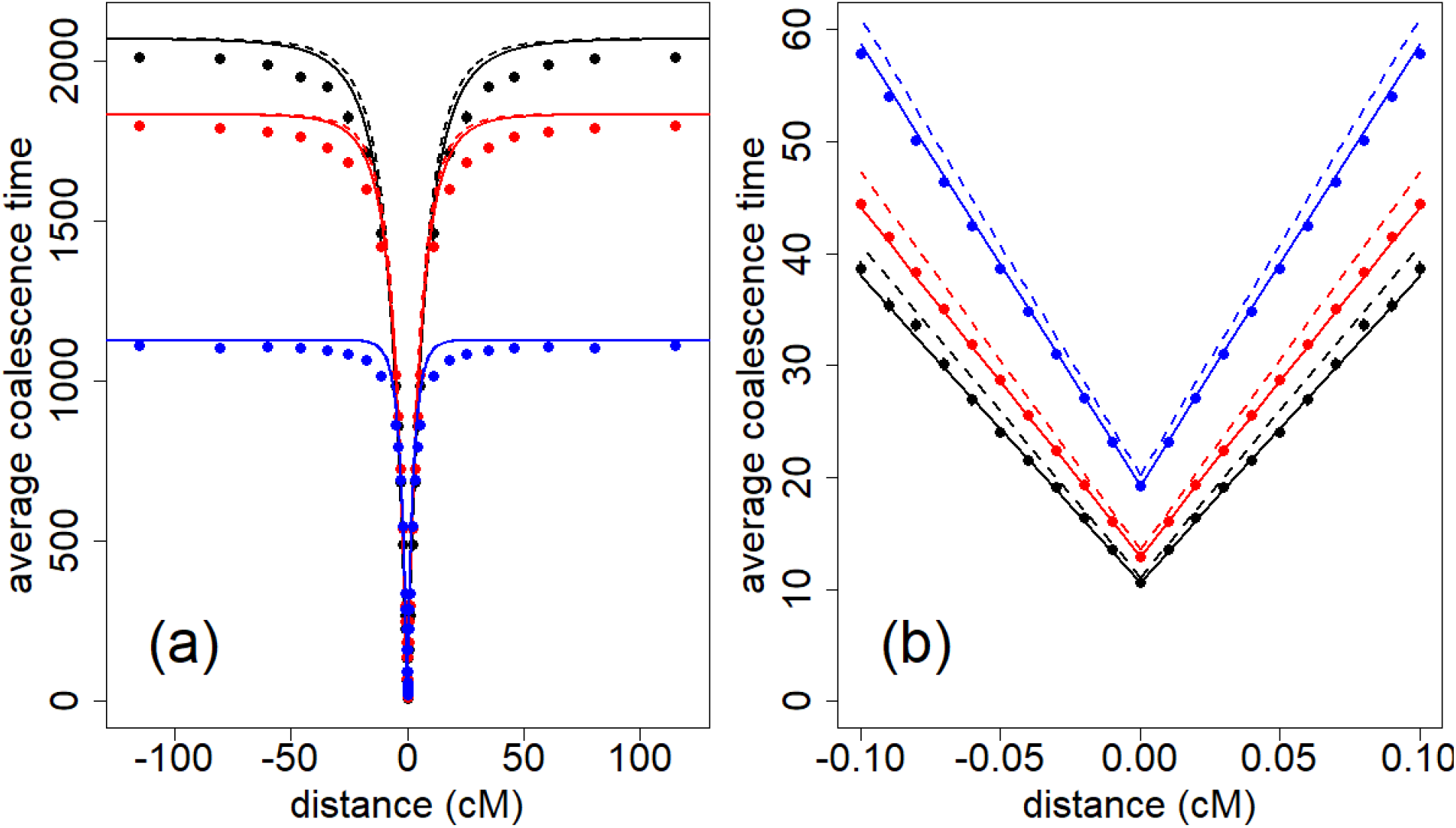
Average coalescence time at a linked locus, as a function of the recombination distance from the focal locus where a mutant fixed in exactly t_m_ generations, starting from a single copy t_m_ generations ago. t_m_ = 15 in black, t_m_ = 20 in red and t_m_ = 40 in blue. The dots are calculated with two-locus WF simulations, and compared to eq. (5) with either a numerical estimation (solid lines) or a theoretical estimation (dashed lines) of 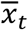 and P(T^(f)^). N_c_ = 20. N_0_ = 1500. The population experienced a contraction t_c_ = t_m_ generations ago.

**Figure 5.**
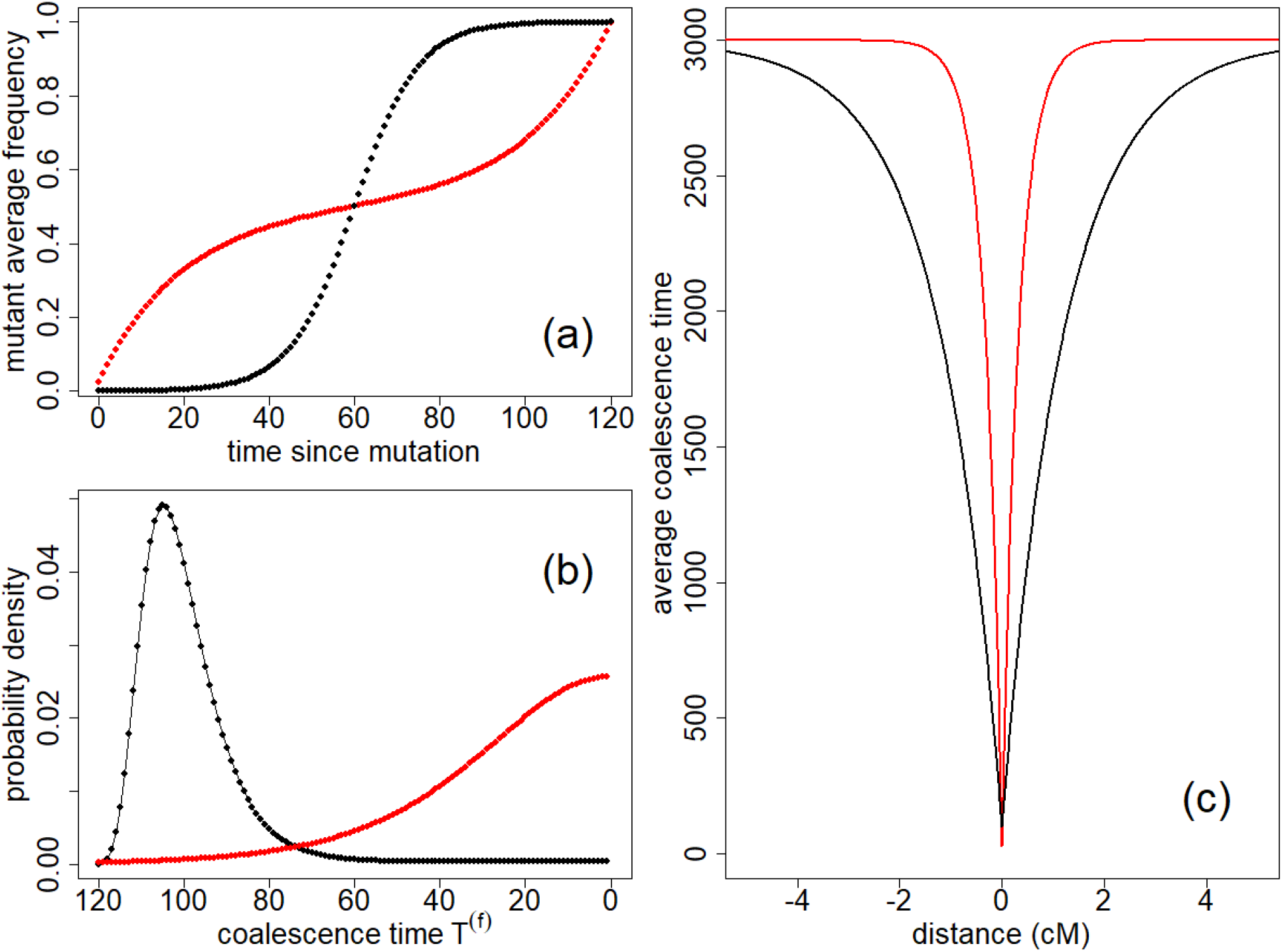
Comparison between troughs of diversity resulting from a selective sweep (black) and a neutral sweep (red), for the same fixation time t_m_ = 120 (corresponding to s ≈ 0.1 in the selective case). Frequency of the fixing allele as a function of time (a), coalescence time distribution (b) and diversity around the fixing site along the genome using eq. (5) (c). N1 = 1500, N_c_ = 20 and N_0_ = 2.97×10^4^.

We observe, as expected, on fig. 4 that the troughs of diversity induced by neutral sweeps are wider and deeper for short fixation times. Similarly to what happens after a selective sweep, there is less opportunity for linked loci to escape the sweep by recombination and maintain diversity when the fixation is fast. In addition, the diversity level at the center of the valley is given by the average coalescence time at the focal locus, which quickly decreases for small fixation times *t*_m_.

### Comparison of neutral sweeps and selective sweeps

Since we did not make any assumption regarding the process driving the mutant allele to fixation when deriving the average coalescence time at linked loci (eq. (2)) and the coalescence time distribution at the focal locus (eq. (3b)), our framework allows us to directly compare the signatures of different processes that can drive mutations to fixation in a given number of generations. We illustrate this by comparing the effect of neutral and hard selective sweeps on linked diversity. Later we will discuss how neutral sweeps compare to a larger variety of scenarios (e.g. background selection, small selection coefficients, or dominant alleles). Here we assume that the neutral and selected fixations occurred over the same time interval, that is in both cases in exactly *t*_m_ generations. The selected fixation is assumed to be codominant (*h*=0.5) and occurs on an autosomal locus in a randomly mating diploid population of constant size *N*1, and we consider a strong selection strength (2*N*_1_s ≫ 1) so that the allele frequency follows the deterministic trajectory

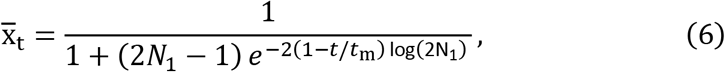

where the fixation time is given by *t*_m_(*s*) = 2log(4*N*_1_*s*)/*s* (Barton 1995). Then combining eqs. (5), (3b) and (6), we can compute the average coalescence time at linked loci as a function of the recombination distance *r* to the focal locus, after replacing *T*_m_, the average coalescence time at *t* = *t*_m_, by 2*N*_1_ in eq. (5) and *N*_c_ by *N*_1_ in eq. (3b). This approach yields results similar to Charlesworth (2020), where the author investigated signals of selective sweeps correcting for coalescent events that happen during the sweep, thus going beyond the common assumption of a star tree structure at the focal locus. For sake of simplicity in the neutral case, we consider that the mutant appeared at the time of the contraction, i.e. *t*_m_ = *t*_c_. Furthermore, we will assume that the average coalescence times (and consequently the genetic diversity) are equal in both scenarios, i.e. that *T*_0_ = 2*N*_1_ which implies that

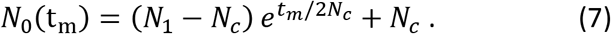

In the neutral case we want the diversity to remain as high as 4*N*_1_ μ after the contraction, which is possible only if the ancestral diversity was even higher, i.e. we have in general *N*_0_ > *N*_1_ > *N*_c_.

In fig. 5a, we compare the mutant average frequency as a function of time for a selected and a neutral fixation. The dynamics of the neutral fixation is the opposite of that of the selected allele in the sense that when one is increasing, the other is “resting” and vice versa. These different trajectories translate into different coalescence distributions at the focal locus (fig. 5b). If selection drives the fixation of the mutation, the distribution of coalescence time is peaked at large coalescence times. In contrast, in the neutral case the distribution is skewed towards small coalescence times. Correspondingly, the coalescence tree for the selected case has a star-like structure (not shown), whereas the tree for the neutral case has shorter outer branches. Therefore, for a given recombination distance, there will be fewer recombinations on the neutral tree because it has a much smaller total length. As recombination helps maintain diversity at linked loci, we would expect neutral troughs of diversity to be wider than in the selected case. However, this is at odds with the valleys of diversity observed in fig. 5c, where the selective trough is wider than the neutral trough. In fact, even though recombination is less abundant on the neutral tree, it is more efficient at recovering diversity. Indeed, if at a linked locus a pair of lineages escapes the sweep due to recombination, it takes on average an extra 2*N*_1_ generations, counted backwards from generation *t* = *t*_m_ when the mutant appeared, for them to coalesce in the selective case, and an extra 2*N*_0_ generations in the neutral case. As *N*_0_ > *N*_1_ two lineages escaping the sweep due to recombination have a larger coalescence time in the neutral case, and correspondingly a larger diversity, which explains why the neutral valley of diversity is narrower. Furthermore, we see that the trough is deeper in the neutral case (fig. 5c), since the average coalescence time is smaller at the focal site due to the smaller total length of the coalescence tree.

To determine if these differences between selective and neutral troughs hold for other fixation times and population sizes, we define two quantities that characterize the shape of a trough, as well as its propensity to be detected in real data: i) the trough relative depth and ii) the width of the trough. The relative depth is defined as the difference between the background level of diversity and the diversity at the focal locus, divided by the background diversity, and the width is measured at half depth, i.e. halfway between the background diversity and the diversity at the focal locus. On fig. 6 we plot the relative depth of neutral and selective troughs as a function of their width for different fixation times *t*_m_, calculated with our analytical expressions. We see that the neutral troughs are not only always narrower than the selective troughs for the same value of *t*_m_, but also deeper. This is due to differences in the focal tree structure between the selective case and the neutral case as well as difference in the ancestral background level in both cases, as explained above. For very short fixation times (corresponding to selection coefficients larger than 0.1), there is almost no difference between troughs generated by selective and neutral sweeps. Indeed, for such values of *t*_m_, in both cases the focal coalescence tree is essentially a star tree because the increase in frequency is very fast, and the ancestral backgrounds of diversity, 2*N*_0_ and 2*N*_1_, are also practically equal. Note however that at small *t*_m_ the corresponding value of the selection coefficient *s* (see legend of fig. 6) may be unrealistically high. For realistic values of the selection coefficient/fixation time, the neutral troughs tend to be quite deep but narrow, whereas selective troughs are wider and their depth decreases quickly for low selection coefficients. From fig. 6, we see that the shape of a neutral trough is generally different from a selective sweep signal, but in practice those differences might be hidden due to the noise inherent present in real genomic data, and it might be difficult to decide whether a genomic signal is a due to a neutral sweep or a selective sweep.

**Figure 6.**
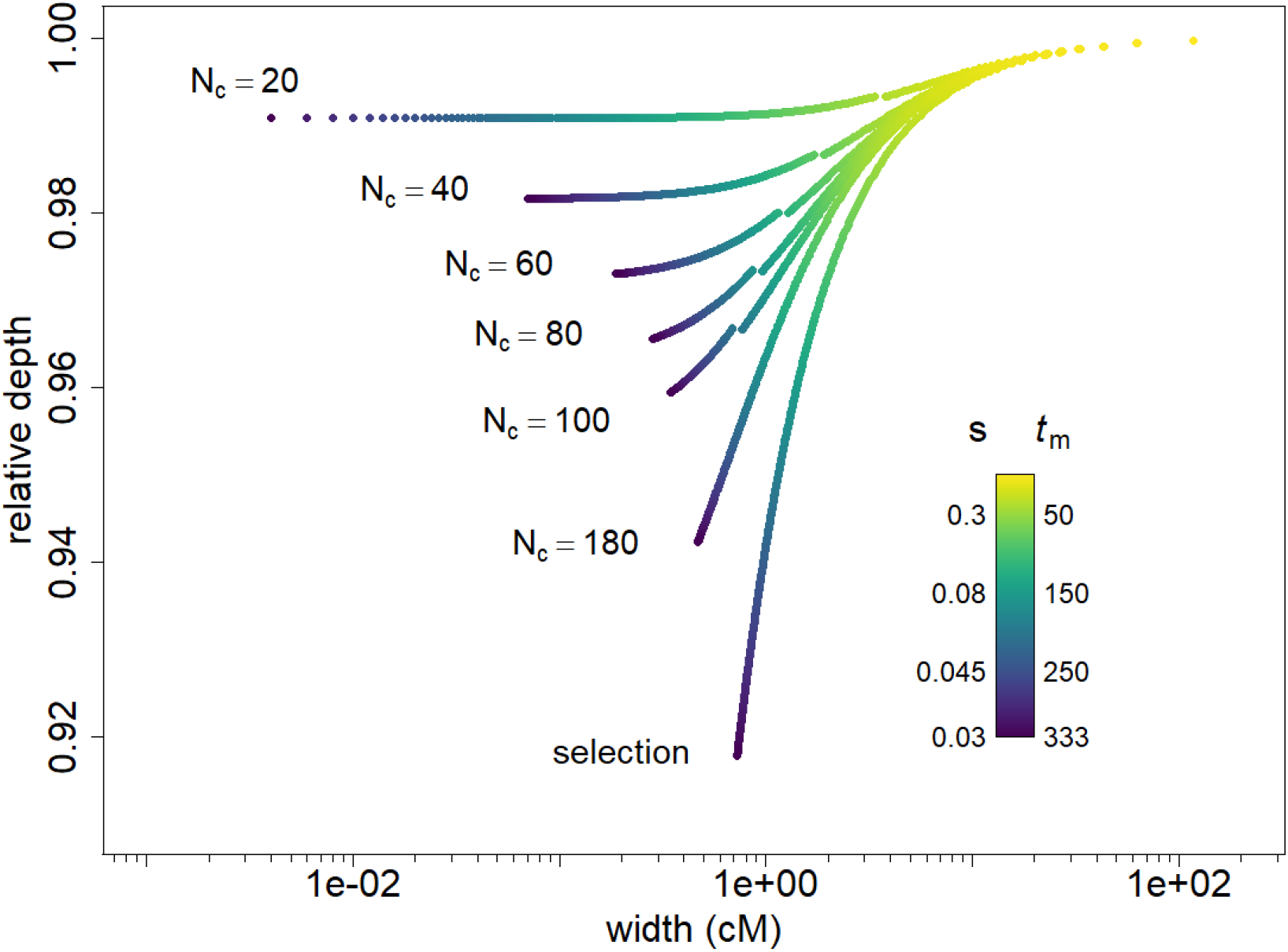
Relative depth as a function of the width of the diversity troughs, for different values of t_m_ and N_c_ in the neutral case and for selective scenarios with identical fixation times. t_m_ goes from 1 to 333 by increments of 1, the corresponding values of the selection coefficient s are indicated on the left of the legend bar (for all of them we have N_1_s ≫ 1). N_1_ = 1500. N_0_ is given by eq. (7) and depends on N_c_ and t_m_. The jumps in the neutral curves for N_c_ = 20, 40, 60, 80 and 100 are due to the use of two different approximations for the frequency of the mutant, eqs. (4a) and (4b) and are located at t_m_ = 2N_c_.

### Is the Qtzl trough in *D. melanogaster* a neutral trough?

A region with reduced nucleotide diversity around the *Quetzalcoatl* gene identified in *Drosophila melanogaster* was judged compatible with a selective sweep (Rogers *et al*. 2010). A hard sweep model (Kaplan *et al*. 1989) was fitted assuming a constant population size of *N*_1_ = 1.85×10^6^ diploid individuals and it was inferred that a positively selected allele fixed in the population 1.5×10^5^ generations ago (1.5×10^4^ years) due to a selective advantage of *s* = 0.0098 (corresponding to a fixation time of more than 300 years). Using our theory, we fitted the data under a neutral demographic scenario of recent population size change that can generate neutral troughs with the same width and almost the same relative depth (less than 0.1% difference) as the *Quetzalcoatl* trough. To infer the demographic parameters, we measure the width of the selective sweep curve used to fit the data in (Rogers *et al*. 2010) and find a set of values of (*N*_c_, *t*_m_) that define a neutral trough with the same width. We then impose that *t*_m_/2*N*_c_ = 0.25 so that the troughs are rare yet observable along the chromosome as explained on fig. 1, and we obtain *t*_m_ = 2200 and *N*_c_ = 4400. In fig. 7 we show a trough generated during a population contraction corresponding to these inferred values, using the software fastsimcoal2 (Excofffier *et al*. 2021). We see that the neutral sweep fit is almost indistinguishable from the selective sweep fit because they not only have the same width, but also practically the same depth. Note that this simulated trough can be also seen in fig. 1c in the region 19-20 Mb. The same approach can be used to generate neutral troughs with a broad range of width and depth, which implies that in most cases, an alternative demographic neutral scenario can be compatible with a trough that is putatively due to selection. In practice, model inference does not rely solely on the fitting of a single trough, and genome wide information must be used. Therefore, we do not exclude here the possibility of the presence of adaptation in the *Quetzalcoatl* gene, but rather make the general warning that valleys of diversity do not necessarily indicate the presence of positive selection.

**Figure 7.**
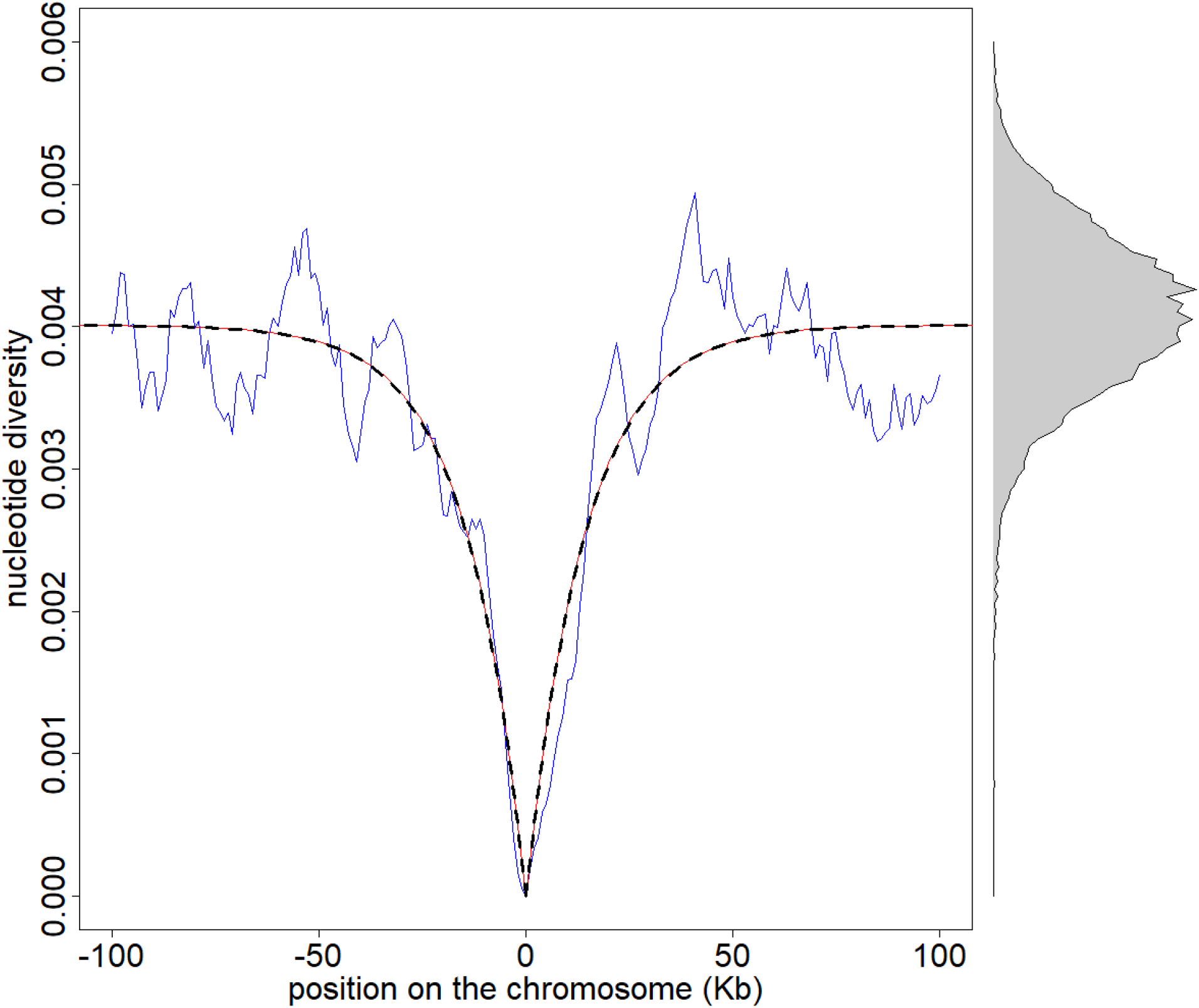
*Trough of nucleotide diversity observed on a 20 Mb chromosome simulated with fastsimcoal2. The population experienced a contraction 2200 generations ago and the (diploid) population size was reduced from N_0_ = 2.37×10^6^ to N_c_ = 4400. The nucleotide diversity (blue line) is calculated on a sample of 30 haplotypes from our simulation. The black dashed line is the expected diversity (eq. (5)) for an allele that just fixed neutrally in the population, starting as a single copy 2200 generations ago. The red line is the expectation of a hard selective sweep with selection coefficient s = 0.0098. On the right we plot the distribution of nucleotide diversity for the whole chromosome. The mutation rate μ = 5.42×10^-10^ per site per generation, and recombination rate r = 3.5×10^-8^ per site per generation were taken from* (Rogers *et al*. 2010). *The nucleotide diversity (in blue) is calculated for sliding windows of 10 Kb at 1 Kb intervals*.

The authors affirm that all data necessary for confirming the conclusions of the article are present within the article, figures, and tables.

## Discussion

It has repeatedly been suggested that strong depletions of diversity in the genome are not necessarily due to the presence of positive selection (Johri *et al*. 2020), and can also be the result of demographic effects only, such as the allele surfing phenomenon occurring at the front of a range expansion (Klopfstein *et al*. 2006). In this work, we considered a model of population contraction to analyze quantitatively the genomic signature of the rapid fixation of a mutation during a population contraction. Taking a step further from previous work that focused on the impact of range expansion on mere allele frequencies, we have studied here the impact of a neutral allele fixation on neighboring genomic diversity. We show that the diversity profile around a recently fixed locus crucially depends on the frequency trajectories of the allele going to fixation, and we outline the fact that neutrally fixing alleles have an inverse-sigmoid trajectory (fig. 3c), as compared to the standard sigmoid frequencies observed for positively selected alleles. For the same fixation time, this difference translates into different genomic signatures (see figs. 5c and 6). Our results demonstrate that there is a short period after a demographic contraction (or during a range expansion) where observed profiles of genomic diversity would look like those usually attributed to selection (fig. 1c), and that selective sweep signals can be mimicked by neutrally fixing mutations without the need to invoke complex histories of population size changes.

Our results allow for a systematic comparison of selective and neutral troughs of diversity, and we used our results to investigate trough shapes for range of neutral and selected scenarios (see fig. 6), which in principle can be used to decide whether a given empirical trough is due to selection or demography, and to infer the corresponding parameters. However, we did not consider the whole spectrum of possible selection scenarios. It would be indeed interesting to use our results to study cases of background selection, small selection coefficients, and a variety of dominance coefficients. All these cases should have their own characteristic trajectories of fixation, and hence potentially different genomic signatures. In addition, in our model we do not consider mutations that fixed in the past (we always assume that the allele has just reached fixation), nor do we consider mutations appearing before the population contraction, i.e., with *t*_m_ > *t*_c_. The average coalescence time in the former case can be expressed as a function of the coalescence time at fixation using conditional probabilities, and we can show that a sweep signal vanishes exponentially with the time elapsed since fixation (see Appendix A4). In the latter case, we can solve the problem by considering the number of gene copies at *t*c that descend from the original copy that appeared at *t*_m_. One could extend our results by considering an allele starting from an arbitrary number of copies at *t*_c_, akin to soft selective sweeps; however, the analytic calculations are complex, and we leave this study for future research. In any case, those additional scenarios must be considered when trying to infer models from the study of troughs found in empirical data. Another phenomenon that renders the inference of parameters cumbersome is a possible interference between troughs. Indeed, when two loci fix neutrally in the population, the genetic diversity in the region between those loci will be influenced by both fixations and will differ from the diversity expected in the vicinity of a single fixing locus. As in the case of interference between the fixation of selected alleles (Weissman and Barton 2012), this should limit the number of independent neutral fixations. The effect of trough interference is stronger for neighboring troughs, and the probability to observe close troughs depends on the relative frequency of troughs along the genome, which itself depends on the distribution of the *T_MRCA_*. In fig. 1d for example, the distribution of *T_MRCA_* has a mode centered around 4*N*_c_ (not shown) and correspondingly the nucleotide diversity is peaked around 4*N*_c_*μ*. As a result, we see many regions of the chromosome with a low diversity. It is likely that those troughs interfere with each other and that they do not correspond to the profile of an isolated trough. On the other hand, in fig. 1c, the first mode of the *T_MRCA_* distribution is truncated because *t*_c_ is much smaller than 4*N*_c_, and only *T_MRCA_*s equal or close to *t*_c_ are observed (plus all the *T_MRCA_*s corresponding to the second mode centered at 4*N*_0_). In this case there is no interference and the (rare) troughs, such as the one in fig. 7, are correctly fitted by their theoretical expectation. Those considerations imply that, even though we know the forward in time probability that an allele will fix in *t*_m_ generations, it is difficult to infer the parameters of a fixation scenario from a single observed neutral valley of diversity. It appears therefore difficult to perform model selection from a single trough signal, i.e., to decide whether a particular trough is due to selection or demographic effects, because alternative demographic scenarios that we did not consider here could also lead to similar signals. In principle, if several troughs of diversity were observed in a genome, one could use the distribution of trough shapes expected under a given simple demographic model and a distribution of fitness effect to compare neutral and selection models under a likelihood framework.

In conclusion, our results suggest that any empirical valley of diversity found in empirical data can be reproduced neutrally with a population contraction using appropriate parameters. One could argue that this identifiability problem disappears once the true evolutionary history is correctly inferred. However, inferring the true demographic history requires precise knowledge about how selection has shaped genomic diversity (Johri *et al*. 2020). In humans, for instance, it has been estimated that roughly 95 % of genomic diversity is affected by some form of non-neutral forces such as background selection or biased gene conversion (Pouyet *et al*. 2018) potentially biasing demographic inference (Ewing and Jensen 2016). These considerations indicate than genome scans in search for signals of adaptation might be subject to stronger false positive rates than previously thought. We thus believe that despite current advances using supervised machine learning or similar approaches (Schrider and Kern 2018), it remains important to further study the effect of neutral fixations in various demographic scenarios using localized genomic approaches such as the present analytical work (Johri *et al*. 2021b); as well as with controlled experiments on real living organisms where both the selected locus and the population history are known (Orozco-terWengel *et al*. 2012). Such work will be critical in order to develop more appropriate evolutionary null models for statistical inference (Hahn 2008; Johri *et al*. 2020).

## Appendix

## A1. Coalescence distribution after a contraction

We want to determine the coalescence time of two lineages in a population that experienced a contraction *t_m_* generations ago, from a diploid size *N*_0_ to *N*_c_. As we go backward in time, the coalescence rate switches from (2*N_c_*)^−1^ to (2*N*_0_)^−1^ at *T* = *t*_c_. The probability distribution might still be approximated by a piecewise exponential density:

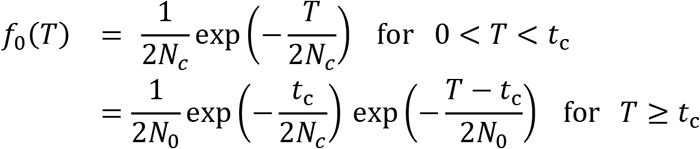

The corresponding expectation for this distribution is

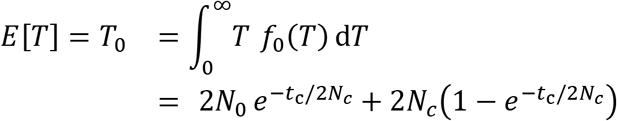

## A2. Average frequency of an allele fixing in exactly *t*_m_ generations

In this section time is counted forward from the mutation, which appears after the contraction, so that during the fixation the diploid population size is constant and equal to *N*_c_. We condition on the fixation time *t*_m_ of the mutant. We define the trajectory of a mutant as the list of frequencies at all generations: {*x_t_*} = (*x*_0_, *x*_1_…, *x*_*t*_m_−1_, *x_t_m__*). We assume that the mutant fixes in exactly *t*_m_ generations, starting from a frequency *p*_0_, i.e. *x*_0_ = *p*_0_, 0 < *x*_*t*_m_−1_ < 1 and *x_t_m__* = 1. The probability that the mutant follows a given trajectory might be expressed as the product of the transition probabilities

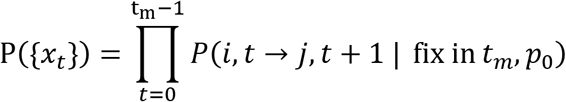

For an unconditional Wright Fisher model, P(i, t → j, t + 1) is the probability to have j copies of the new allele at t + 1 given that there were i copies at t. We note *P_t_*(*i* → *j*) for brevity. If we only consider trajectories fixing in exactly *t_m_* generations and starting from a number 2*N_C_ p*_0_ of copies at t = 0, then the transition probabilities are not equal to the transitions of the unconditional Wright-Fisher model. However, thanks to Bayes theorem, we can write

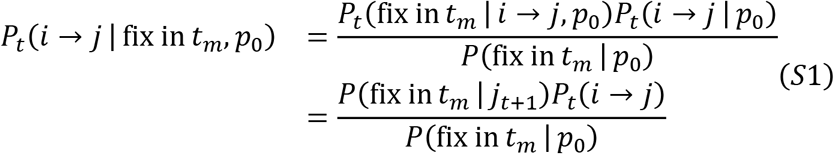

From the first to the second line, we use the Markov property. The three terms involved in the right-hand side of this equation can be approximated thanks to diffusion theory. In this framework, the probability for an allele to fix in *t_m_* generations, given that there were i copies at time t is approximately (Ewens 2004, taking the time derivative of eq. 5.39)

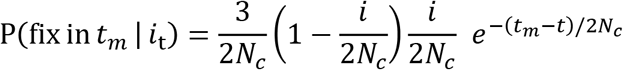

The term *P_t_*(*i* → *j*) is the unconditional binomial transition probability of the Wright Fisher model (which does not depend on t). In principle, eq. (S1) can be used to compute the exact distribution of coalescence times at the focal locus, using eq. (3a). However, the huge number of possible trajectories fixing in *t_m_* generations ((2*N_c_* – 1)^t_m_−1^) makes the average over trajectories impossible to evaluate numerically. For this reason, we use the approximation in eq. (3b).

We consider here the probability that the allele has frequency x at time t, given that it started at frequency *p*_0_ at t = 0. Again if we only consider trajectories that fix in exactly *t_m_* generations, this probability is not equal to the neutral diffusive result. However, similarly to the previous section, we can use Bayes theorem:

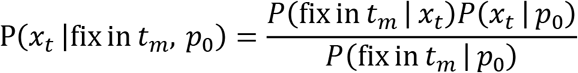

From diffusion theory (Ewens 2004, eq. 5.11), we also have

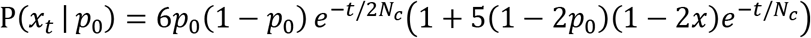

which is a second order expansion of an infinite series involving vanishing exponential terms (*e*^−*k*(*k*+1)*t*/4*N_c_*^ for all k ≥ 1). This expansion is thus valid in the limit of large times t ≥ 2*N_c_*. We deduce that the probability that an allele fixing in *t_m_* generations has frequency x at time t is

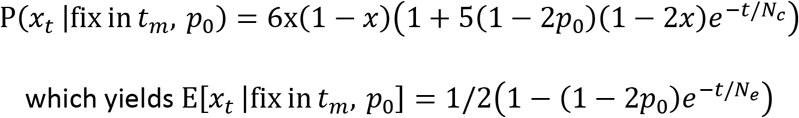

This expression is valid for t_m_ ≫ t ≫ 2*N_c_*, and does not allow to estimate the frequency close to fixation (we see that E[*x_t_*] tends to 1/2 as time grows). However, invoking a symmetry argument we may write

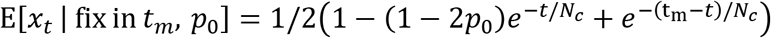

When t_m_ ≪ 2*N_c_*, we can use a linear approximation for the trajectory (based on the numerical observations)

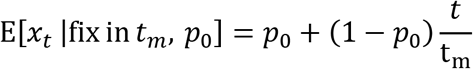

## A3. Coalescence distribution at linked loci around a neutral fixation

We now return to the scenario of fig. 2, with a backward in time approach. Using Bayes theorem, we express the coalescence time of two haplotypes at the linked locus *T*^(*l*)^ conditioning on the coalescence time at the focal locus *T*^(*f*)^

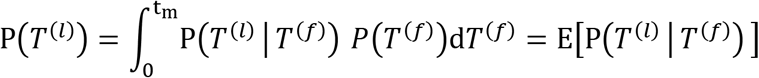

We assume that the linked locus is close to the focal locus on the chromosome, more precisely that the recombination rate r is very small r ≪ 1, so that we consider at most one recombination, occurring on one of the two focal lineages. We distinguish cases where there is no recombination between *t* = 0 and *t* = *T*^(*f*)^, cases where the allele at the linked locus recombines (somewhere between *t* = 0 and *t* = *T*^(*f*)^) onto a haplotype carrying the ancestral allele at the focal locus, and cases where the allele at the linked locus recombines onto a haplotype carrying the derived allele at the focal locus. We call the second and third case homozygous and heterozygous recombination respectively, referring to the zygosity at the focal locus of the recombining pair of haplotypes (note that are three haplotypes, the two first ones have a coalescence time *T*^(f)^, and the third one recombines with one of these two). If there is no recombination, then the coalescence time is the same for both loci, *T*^(l)^ = *T*^(f)^. To treat the case with a homozygous recombination, it is convenient to name the haplotypes: *i* and *j* coalesce at *T_ij_*^(f)^ = *T*^(f)^ at the focal locus, and *k* is a third haplotype, onto which the linked allele recombines (coming from *i*). The linked allele carried by *j* stays on the same haplotype (no more than one recombination), and after recombining onto *k*, the linked allele initially carried by *i* also stays on *k* (again, at most one recombination). This implies that those two linked alleles coalesce at *T_ij_*^(l)^ = *T_jk_*^(f)^. This time is in general different than *T_ij_*^(f)^, however on average *T_jk_*^(f)^ tand *T_ij_*^(f)^ are equal (averaging over all possible coalescence trees at the focal locus). This implies that we can treat the case with homozygous recombination as if there was no recombination. If there is a heterozygous recombination between *i* and *k*, at some generation between *t* = 0 and *t* = *T*^(f)^, then the linked alleles still have not coalesced at *t* = *t_m_* because after the recombination one of them is linked to a derived focal allele and the other one to an ancestral focal allele (and they stay linked because there is at most one recombination). In that case, *T_ij_*^(l)^ is equal to *t_m_* plus a random time given by (on average) *T_m_*, and is independent of *T^ij^*^(f)^. Using again Bayes theorem and the previous results to write

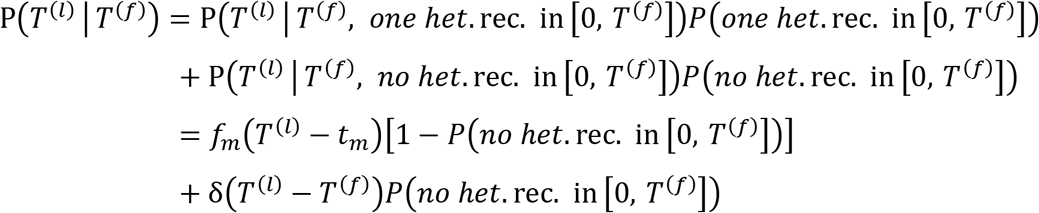

Where *δ*(·) is the Dirac delta function, and *f_m_* is the unconditional coalescence distribution of a pair of lineages sampled at *t* = *t_m_*, i.e. it is equal to the function *f*_0_ introduced above but replacing *t_c_* by *t_c_* – *t_m_* (note also that *f_m_*(*t*) = 0 if *t* < 0). We then have to evaluate the probability that there is no heterozygous recombination. At generation *t* (counted backward) the probability that a linked allele recombines onto a haplotype carrying the ancestral allele at the focal locus is *r*(1 – *x_t_*), where *x_t_* is the frequency of the derived allele at the focal locus, we deduce that the probability that there is no heterozygous recombination on either lineage is

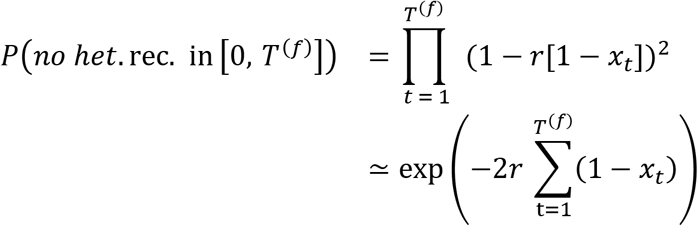

This probability depends explicitly on the allele trajectory, which means that rigorously, all the calculations should be conditioned on a given trajectory, and then averaged over all trajectories. To allow for mathematical tractability, and to avoid heavy expressions, we consider that as a good approximation 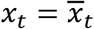. Finally we obtain

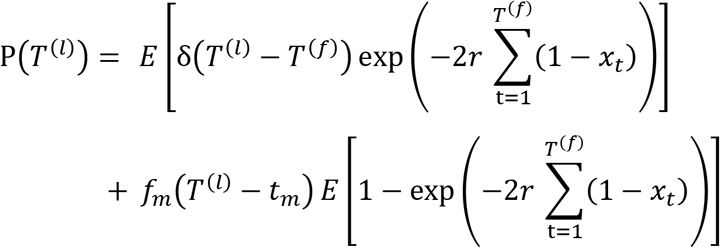

The expectation corresponding to this distribution yields eq. (2).

## A4. Average coalescence time at a linked locus around a mutation that completed fixation *t*_fix_ generations ago

Thanks to Bayes theorem we can write

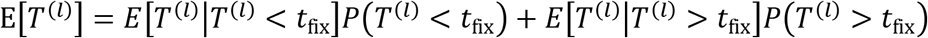

i.e. we distinguish coalescence events happening in less than *t*_fix_ generations or more than *t*_fix_ generations. In the former case, the coalescence is neutral, unconditional (the fixation is completed) and happens in a population of constant size *N*_c_ which means that *E*[*T*^(*l*)^|*T*^(*l*)^ < *t*_fix_] and *P*(*T*^(*l*)^ < *t*_fix_) can be worked out from the neutral exponential distribution. On the other hand, *E*[*T*^(*l*)^|*T*^(*l*)^ > *t*_fix_] is equal to *t*_fix_ plus the expectation from eq. (5) which we note here E[*T*^(*l*)^] (*t* = *t*_fix_). We obtain

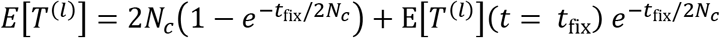

We see that the sweep signal vanishes exponentially with the time elapsed since fixation.

## Acknowledgment

This work was partially supported by a Swiss NSF grant No 310030_188883 to LE. We are grateful to Montgomery Slatkin, Brian Charlesworth and Jeff Jensen for their helpful comments.

